# Tumor protein D54 binds intracellular nanovesicles *via* an amphipathic lipid packing sensor (ALPS) motif

**DOI:** 10.1101/2021.12.03.471088

**Authors:** Antoine Reynaud, Maud Magdeleine, Amanda Patel, Anne Sophie Gay, Delphine Debayle, Sophie Abelanet, Bruno Antonny

## Abstract

Tumor Protein D54 (TPD54) is an abundant cytosolic protein that belongs to the TPD52 family, a family of four proteins (TPD52, 53, 54 and 55) that are overexpressed in several cancer cells. Even though the functions of these proteins remain elusive, recent investigations indicate that TPD54 binds to very small cytosolic vesicles with a diameter of ca. 30 nm, half the size of classical transport vesicles (e.g. COPI and COPII). Here, we investigated the mechanism of intracellular nanovesicle capture by TPD54. Bioinformatical analysis suggests that TPD54 contains a small coiled-coil followed by several amphipathic helices, which could fold upon binding to lipid membranes. One of these helices has the physicochemical features of an Amphipathic Lipid Packing Sensor (ALPS) motif, which, in other proteins, enables membrane binding in a curvature-dependent manner. Limited proteolysis, CD spectroscopy, tryptophan fluorescence and cysteine mutagenesis coupled to covalent binding of a membrane sensitive probe show that binding of TPD54 to small liposomes is accompanied by large structural changes in the amphipathic helix region. TPD54 binding to artificial liposomes is very sensitive to liposome size and to lipid unsaturation but is poorly dependent on lipid charge. Cellular investigations confirmed the key role of the ALPS motif in vesicle targeting. Surprisingly, the vesicles selected by TPD54 poorly overlap with those captured by the golgin GMAP-210, a long vesicle tether at the Golgi apparatus, which displays a dimeric coiled-coil architecture and an N-terminal ALPS motif. We propose that TPD54 recognizes nanovesicles through a combination of ALPS-dependent and -independent mechanisms.

## Introduction

In cells, transport vesicles are produced from membrane-bound compartments by vesicular coat complexes, which self-assemble at the membrane surface into spherical shells to promote the budding of coated-vesicles (1, 2). The best characterized protein coats are COPII, COPI, AP1-clathrin and AP2-clathrin, which promote vesicle budding from the endoplasmic reticulum (ER), Golgi apparatus (GA), trans Golgi network (TGN), and plasma membrane (PM), respectively. These different coats and the corresponding classes of vesicles have been abundantly studied over the last decades, notably as regards to the mechanisms of coat assembly, coat structure, vesicular content in cargoes, and associated diseases (3, 4). Investigations by electron microscopy both *in situ* and from reconstituted systems indicate that these vesicles have a well-defined diameter, which is in the range of 80 nm (5, 6). Although protein coats can adapt their size to form bigger vesicles for the packaging of large cargoes (7, 8), the converse has not been observed and transport vesicles smaller than 50 nm have rarely been reported.

Surprisingly, a new class of transport vesicles has been recently identified through cellular investigations of Tumor Protein D54 (TPD54) (9). TPD54 is a small (206 aa, 22.2 kDa) protein that belongs to a family of four proteins: TPD52, TPD53 (or TPD52L1), TPD54 (or TPD52L2) and TPD55 (or TPD52L3) (10). These proteins are ubiquitously expressed in cells except TPD55, which is specific to testis. Light microscopy imaging indicates that TPD54 associates to various membrane-bound compartments, mostly Golgi apparatus and endosomal structures (9). In addition, a large fraction of TPD54 is found in the cytosol but this localization is deceptive. Fluorescence Recovery After Photo Bleaching (FRAP) experiments suggests that the cytosolic fraction of TPD54 corresponds to very small structures (≈ 32 nm), which could be resolved by super resolution microscopy (9). When TPD54 is redirected to the mitochondria by a knocksideway-based approach, it causes the accumulation of numerous small-sized vesicles (diameter ≈ 30 nm) at the mitochondria surface. Several lines of evidence suggest that these vesicles are involved in membrane traffic. First, deletion of TPD54 causes a severe Golgi dispersal phenotype as well as a delay in the secretion of a model cargo (E-cadherin). Second, TPD54 vesicles contain many Rab proteins, which are small G proteins involved in vesicular traffic, as well as Q-SNARE proteins, which promote vesicle fusion (9).

Here, we show that TPD54 recognizes highly curved membranes through the folding of an amphipathic lipid packing sensor (ALPS) motif. ALPS are amphipathic sequences of 20 to 40 aa motifs that are characterized by the abundance of small polar residues (Gly, Ser, Thr) and by the presence of regularly spaced bulky hydrophobic residues (11-15). ALPS motifs are intrinsically unstructured but fold as amphipathic helices at the surface of highly curved membranes displaying lipid packing defects (13, 16-19). ALPS motifs are present in several proteins that transiently interact with highly curved membranes (20). These include the COPI coat regulators ArfGAP1 and Gcs1 (15, 21-24), and vesicle tethering factors such as the golgin GMAP-210 (13, 14, 25, 26), the HOPS complex (27, 28) and synapsin (29). In addition, functional ALPS motifs are present in the sterol transfer protein Osh4 (30, 31), the autophagy protein ATG14L (32-34), the Golgi-associated PI-4 kinase PI4KIIIß (35) and some members of the nuclear pore complex (13, 36).

We show that TPD54 shows a sharp preference for highly curved and unsaturated lipid membranes *in vitro* and, as such, resembles previously characterized ALPS-containing proteins. In cells, mutating the ALPS motif of TPD54 abolished its association to disparate membranes. However, differences in localization between TPD54 and the golgin GMAP-210, which also captures small transport vesicles through an ALPS motif, suggest the involvement of additional interactions for the selection of specific small transport vesicles. In line with this hypothesis, binding of TPD54 to small liposomes is accompanied by a large structural change that applies not only to the ALPS motif but also to the full region downstream of the coiled-coil region.

## Results

### Bioinformatic analysis of TPD54

**Figure 1**. shows the domain organization of human TPD54 (206 aa) as deduced from several bioinformatic tools and previous experiments (9). The PONDR algorithm (http://www.pondr.com/), which predicts the probability of a sequence to contain intrinsically disordered regions, suggests that the protein is largely unstructured (**Figure 1A**). However, TPD54 contains a predicted coiled-coil region spanning from residue 40 to 80 (https://embnet.vital-it.ch/software/COILS_form.html) (**Figure 1B**). This region has been shown to contribute to homo and heterotypic associations between TPD protein members (37). However, the exact geometry (parallel *vs* antiparallel) and stoichiometry (dimer, trimer or tetramer) of the coiled-coil is not known. The recently published machine learning method Alphafold (38) (https://alphafold.ebi.ac.uk/) overall agrees with this architecture, predicting with high confidence (> 0.9) an α-helix between aa E43 and R79 corresponding to the coiled-coil, whereas other regions are predicted as either random coils (M1-T42; V178-F206) or α-helical (T85-V102; S104-Q122; D124-R159; A162-K177) but with low or medium confidence (**Figure 1C**).

**Figure 1.**
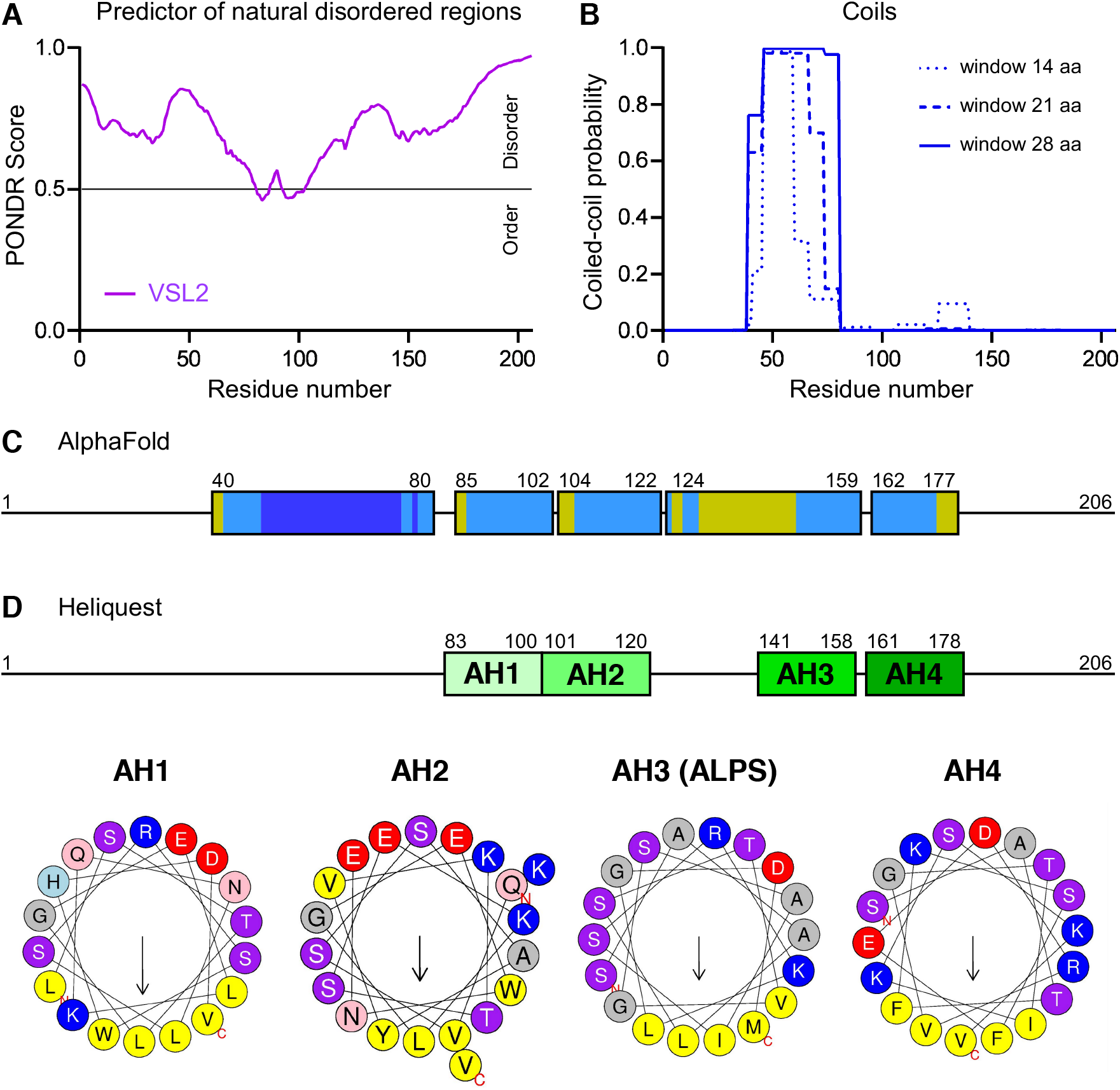
Bioinformatic analysis of TPD54. **A**. *PONDR* analysis of the TPD54 sequence. **B**. Coiled-coil prediction with *Coils*. **C**. Helical structures as predicted by *AlphaFold*. The color code shows the degree of confidence: high, dark blue; medium, light blue; low, yellow. **D**. Helical projections of the indicated sequences of TPD54 using *Heliquest*. The analysis suggests the presence of four amphipathic helices. Color code: yellow, hydrophobic residues; blue, basic; red, acidic; purple, Ser and Thr; grey, Ala and Gly; pink, Gln and Asn.

We previously used the bioinformatic tool HELIQUEST (https://heliquest.ipmc.cnrs.fr/) (39) to detect the presence of ALPS motifs in the full human and yeast proteomes and identified TPD54 as one of the ≈ 200 human proteins containing an ALPS motif (13). The ALPS motif of TPD54 spans S141-M158. By rescanning the full TPD54 sequence with HELIQUEST, we observed that three other regions, two upstream and one downstream of the ALPS motif, also display an amphipathic helical sequence (**Figure 1D**). They are referred to as AH1 (L83-V100), AH2 (Q101-V120) and AH4 (T163-G180), with AH3 corresponding to the ALPS motif. They all contain Gly residues, which should make them malleable for switching from a random coil to an α-helix in the presence of a proper membrane template. AH1, AH2 and AH4 are rich in both positively and negatively charged residues in contrast to ALPS motifs (13, 15). Of note, AH1-AH2 as well as AH3-AH4 are not in register, which means that if they were forming a single continuous α-helix, their respective hydrophobic faces would point toward opposite directions.

Overall, the predicted domain organization of TPD54 with both a coiled-coil region and amphipathic motifs bears resemblance to the N-terminal region of the golgin GMAP-210, a very long coiled-coil protein. GMAP-210 acts as a tethering string, capturing small transport vesicles through its N-ter ALPS motif (14, 25, 40).

### TPD54 binds to simple phospholipid vesicles in a curvature-dependent manner

We purified recombinant human TPD54 from a GST fusion form by a three-step protocol: (i) glutathione-bead binding step; (ii) thrombin cleavage of the linker between GST and TPD54; (iii) size exclusion chromatography. To determine if TPD54 can directly interact with lipid membranes, we incubated full-length TPD54 with liposomes and recovered the liposome-bound fraction by flotation after centrifugation on sucrose cushions. **Figure 2A** shows the result of an experiment with three preparations of liposomes: PC(18:1/18:1) (dioleoyl-phosphatidylcholine) obtained extrusion through 200 nm polycarbonate filters; PC(18:1/18:1) obtained by sonication; and PC(4ME 16:0/4ME 16:0) (diphytanoyl-PC) obtained extrusion through 200 nm polycarbonate filters. Diphytanoyl-lipids, which are not present in eukaryotic cells, form lipid membranes that are very permissive to the binding of amphipathic helices owing to the branched nature of their acyl chains, which favors hydrophobic insertions (41). TPD54 bound weakly to 200 nm PC(18:1/18:1) liposomes, but strongly to sonicated PC(18:1/18:1) liposomes and to 200 nm PC(4ME 16:0/4ME 16:0) liposomes. Thus, TPD54 is able to interact directly with simple membranes (i.e. PC) and binding is very sensitive to both membrane curvature and lipid acyl chain composition.

**Figure 2.**
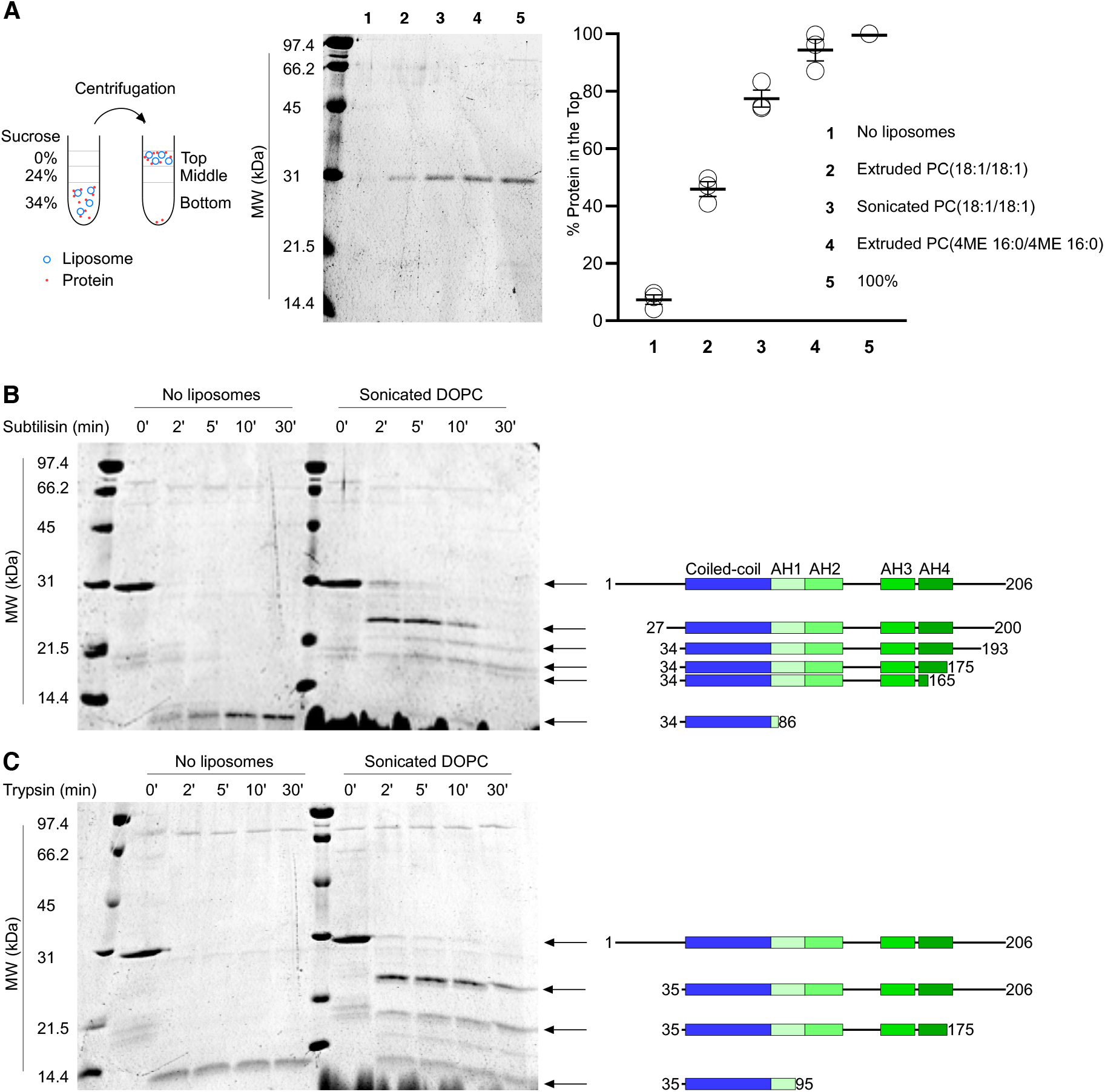
Limited proteolysis indicates that binding of TPD54 to small liposomes occurs through a large conformational change. **A**. Flotation experiments of TPD54 with liposomes. After incubation of purified TPD54 with liposomes, the liposomes were recovered by flotation on sucrose cushions and analyzed by SDS-PAGE using Sypro orange staining. The open circles in the plot show the results of three independent experiments. The bars show the mean ± SEM. **B, C**. limited proteolysis of TPD54 with trypsin or subtilisin in the presence or in the absence of sonicated PC(18:1/18:1) liposomes. The sequence of the various fragments was determined by mass spectrometry.

### Large structural changes accompany the binding of TPD54 to membranes

To determine if structural changes accompany the binding of TPD54 to membranes, we conducted limited proteolysis experiments. For this aim, we incubated TPD54 with trypsin or subtilisin and analyzed the resulting fragments over time by SDS-PAGE and mass spectrometry. Trypsin cleaves peptide bonds downstream of accessible Lys or Arg residues, whereas subtilisin cleaves accessible peptide bonds with poor sequence specificity. We noticed that TPD54 showed anomalous migration on SDS-PAGE with an apparent MW close to 30 kDa, much higher than the calculated MW (22368.32 Da) and the observed MW by mass spectrometry (22367.245 Da) (**Figure 2B**). This anomalous migration was systematic for all fragments analyzed and might be related to weaker SDS binding due to some intrinsic features of the TPD54 sequence (42).

In the absence of liposomes, trypsin and subtilisin rapidly digested TPD54, leading to the apparition of a single protease-resistant band with an apparent size below the lowest MW marker (14 kDa; **Figure 2B**). Mass spectrometry identified this band as aa 34-89 and aa 35-90 in the case of subtilisin and trypsin digestion, respectively. In agreement with the bioinformatic analysis (**Figure 1**), this result suggests that only a small part of TPD54, which corresponds to the predicted coiled-coil region, is intrinsically folded, whereas the remaining regions display no stable structure.

Incubation of TPD54 with sonicated liposomes caused an important protection of TPD54 from protease degradation. For both subtilisin and trypsin, a fragment with an apparent MW ≈ 24 kDa rapidly accumulated at t = 2 to 5 min before subsequent digestion into smaller fragments of apparent MW of 23 to 19 kDa (**Figure 2B**). Analysis by mass spectrometry indicate that the first 26-34 amino-acids, which are upstream of the predicted coiled-coil, were rapidly degraded. In contrast, the region downstream of the coiled-coil region appeared essentially intact at early time points (t = 2 to 5 min) and became slowly digested into smaller fragments according to the series aa 206 > 200 > 193 > 175 > 165 (i.e. up to the fourth predicted AH) (**Figure 2B**). These results suggest that the recognition of small vesicles by TPD54 leads to major changes in the protein structure, which applies to the full region downstream of the predicted coiled-coil region.

### TPD54 binding to small liposomes is accompanied by α-helical folding

Given the fact that the region that becomes protected from proteolysis in the presence of small liposomes contains the four predicted amphipathic helices (AH1-AH4), we performed circular dichroism experiments to determine if changes in the secondary structure accompanies TPD54 adsorption to small liposomes. In solution, the CD spectrum of TPD54 featured two minima at 208 and 222 nm, which are characteristic of α-helical structures. Upon addition of sonicated PC(18:1/18:1) liposomes, these two negative peaks further increased, suggesting additional folding of α-helical regions (**Figure 3A**). Fitting the CD spectra using the web tool CAPITO (43) suggests that the content in α-helix increased from 30% in solution to 60% in the presence of sonicated PC(18:1/18:1) liposomes. Considering that the coiled-coil region of TPD54 is 45 aa whereas each predicted amphipathic helix (AH1 to AH4) is about 18 aa in length, this two-fold increase in helicity suggests the folding of the equivalent of 2 to 3 of these helices.

**Figure 3.**
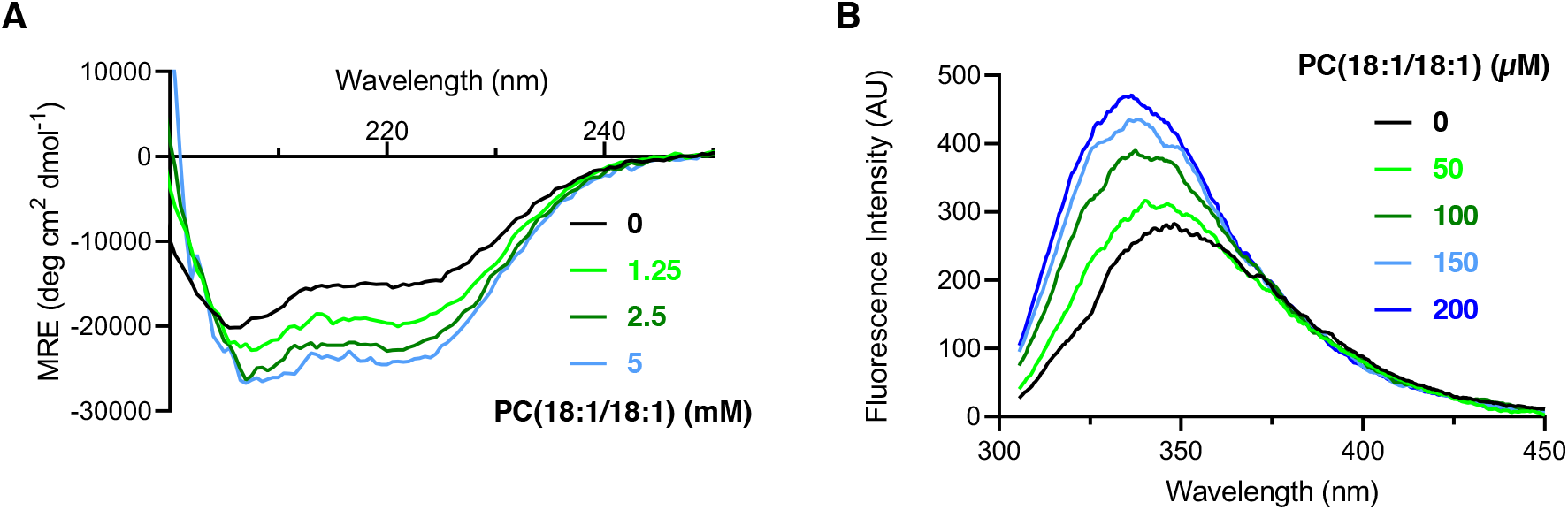
Binding of TPD54 to small liposomes induces changes in circular dichroism and Tryptophan fluorescence. **A**. CD spectrum of TPD54 in the absence or in the presence of increasing concentration of sonicated PC(18:1/18,1) liposomes. **B**. Fluorescence emission spectrum of TPD54 in the presence of increasing concentrations of sonicated PC(18:1/18,1) liposomes.

### Tryptophan fluorescence reveals conformational changes in the 85-140 region

The TPD54 region downstream of the predicted coiled-coil contains two tryptophan residues (W97 and W116) whereas no other tryptophan is present in the TPD54 sequence (**Figure 1D**). Because tryptophan fluorescence is very sensitive to environmental changes and because this region becomes fully protected from protease degradation in the presence of sonicated PC(18:1/18:1) liposomes, we compared the intrinsic fluorescence of TPD54 in the absence and in the presence of such liposomes. As shown in **Figure 3B**, binding of TPD54 to sonicated PC(18:1/18:1) liposomes promoted a large tryptophan fluorescence increase, which was accompanied by a blue shift. This observation further strengthens the hypothesis that binding of TPD54 to curved membranes leads to a large reorganization of the protein structure, including the region encompassing the two tryptophan groups.

According to *Heliquest* (39), W97 and W116 belong to the hydrophobic faces of the putative amphipathic helices AH1 and AH2, respectively (**Figure 1D**). To determine whether tryptophan W97 and/or W116 become close to the membrane interfacial region upon TPD54 binding to small liposomes, we conducted fluorescence resonance energy transfer (FRET) experiments. We used PC(18:1/18:1) liposomes doped with 5 mol% of the fluorescent lipid diphenylhexatriene PC (DPH-PC). The excitation spectrum of DPH-PC overlaps with the emission spectrum of tryptophan, thereby leading to a FRET signal when tryptophan groups partition in lipid membranes (44, 45). Stepwise additions of TPD54 to sonicated DPH-PC - containing liposomes induced a FRET signal, whereas no increase was observed with extruded liposomes (**supplementary information - Fig S1**). These results suggest that W97 and/or W116 become close enough to the membrane interface (FRET distance is typically in the nm range) when TPD54 adsorbs onto highly curved PC liposomes.

### Membrane proximity analysis using NBD-cysteine scanning mutageneis

To map the TPD54 membrane-interacting regions in a more extensive manner, we prepared various cysteine mutants, which we labelled with the membrane-sensitive fluorescent probe NBD. We selected seven positions, which flank the five predicted structural elements in the following order: A37C - coiled-coil - G82C - AH1 - Q101C - AH2 - S123C - G137C - AH3 - S161C - AH4 - G185C (**Figure 4**). In flotation experiments, all mutants showed stronger binding to sonicated PC(18:1/18:1) liposomes than to extruded ones (200 nm) (**supplementary information - Fig S2**). However, we observed strong differences between the NBD-labelled forms when we recorded their fluorescence in the presence or absence of liposomes. When NBD flanked the predicted amphipathic helices AH1, AH2 and AH3 (mutants G82C-NBD, Q101C-NBD, S123C-NBD, G137C-NBD and S161C-NBD), we observed a large (2 to 3-fold) increase in NBD fluorescence upon the addition of sonicated PC(18:1/18:1) liposomes or extruded (200 nm) PC(4ME 16:0/4ME 16:0) liposomes (**Figure 4A, red and blue spectra, respectively**). As expected, no fluorescence change was observed in the presence of large PC(18:1/18:1) liposomes (**Figure 4A, green spectra**). In contrast, TPD54 A37C-NBD, in which the fluorescent probe is upstream of the predicted coiled-coil, showed no fluorescence change, whereas TPD54 G185C-NBD, in which the fluorescent probe is downstream of the fourth predicted amphipathic helix (AH4) showed a modest (x 1.2 – 1.3) fluorescence increase in the presence of either sonicated DOPC liposomes or extruded (200 nm) PC(4ME 16:0/4ME 16:0) liposomes. Overall, the results of the NBD cysteine scanning mutagenesis, which are summarized in **Figure 4B**, were in good agreement with the limited proteolysis experiments and suggested that the three predicted amphipathic helices AH1, AH2 and AH3 become close to the membrane interface when TPD54 adsorbs to membranes.

**Figure 4.**
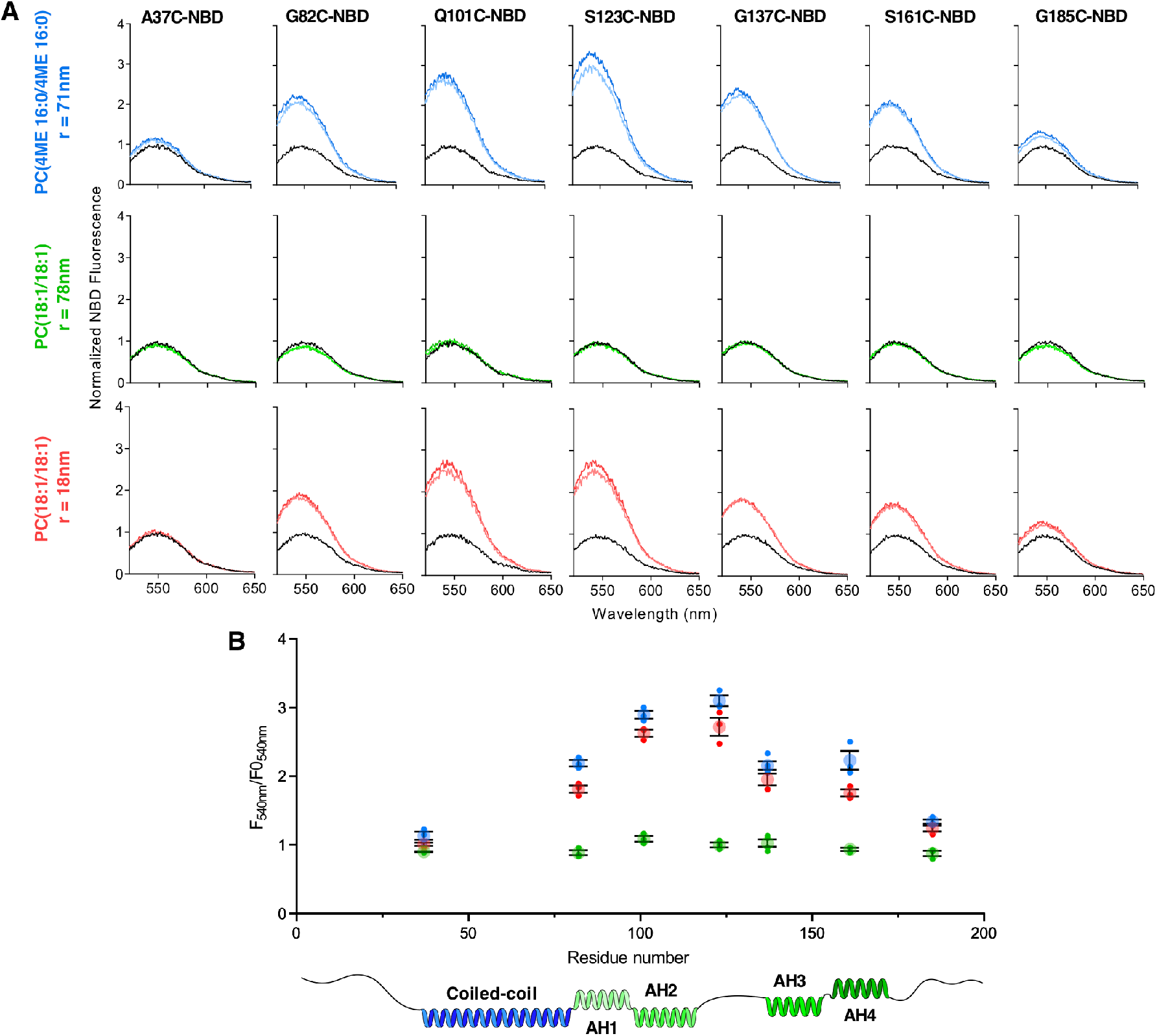
Mapping the TPD54 membrane interaction by NBD fluorescence. **A**. Emission fluorescence spectra of NBD-labelled TPD54 (100 nM) at the indicated position in buffer (black curves) or in the presence of 50 µM (light color curves) or 100 µM PC liposomes (dark color curves). The liposomes were extruded (200 nm) PC(4ME 16:0/4ME 16:0) (blue curves), extruded (200 nm) PC(18:1/18:1) (green curves) or sonicated PC(18:1/18:1) (red curves). **B**. Quantification of the change in NBD fluorescence at 540 nm as a function of the position of the labelled cysteine from three independent experiments similar to that shown in A. The small symbols show the results of each experiment. The large symbols and associated error bars show mean ± SEM.

### TPD54 binds cellular structures through its ALPS motif

Next, we tested the contribution of AH1, AH2, AH3 (ALPS) and AH4 to the subcellular distribution of TPD54. For this aim, we expressed full-length TPD54, which was N-terminally fused with GFP, in RPE1 cells and analyzed the effects of mutations that should decrease the interaction of these putative amphipathic helices with the lipid membrane. We selected four mutations L93D (AH1), L113D (AH2), I151D (AH3), and V171D (AH4), which introduce a negative charge in the hydrophobic face of the putative helices.

At moderate expression level, wild-type GFP-TPD54 showed a composite distribution pattern. In agreement with a previous study (9), GFP-TPD54 distributed between a cytosolic pool, a perinuclear fraction and some rounded structures scattered in the cytoplasm (**Figure 5A**). The perinuclear fraction corresponded to the Golgi apparatus, showing apposed localization with Golgi markers (**supplementary information - Fig S3**). Mutating the first (L93D) or the fourth (V171D) predicted amphipathic helix caused no significant change in the distribution of the protein, whereas mutating the second (L113D) predicted amphipathic helix slightly reduced the Golgi pool of TPD54 (**Figure 5A, B**). Strikingly, when we introduced the disruptive I151D mutation in AH3, that is the ALPS motif, GFP-TPD54 no longer decorated intracellular structures and appeared essentially cytosolic (**Figure 5A, B**).

**Figure 5.**
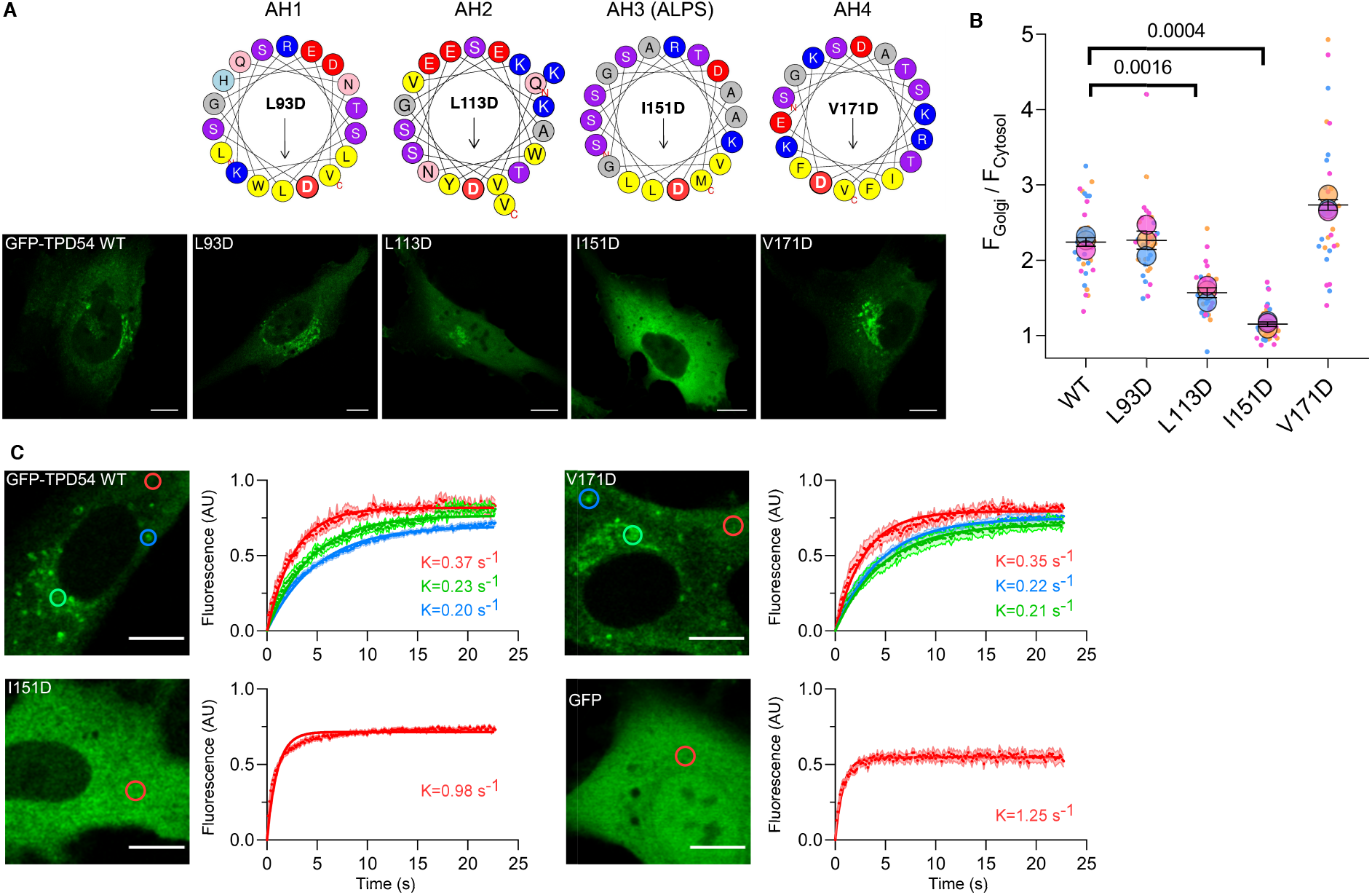
The ALPS motif contributes to TPD54 subcellular localization. **A**. Representative confocal images of RPE1 cells expressing wild-type GFP-TPD54 or mutants harboring a negative charge in one of the four putative amphipathic helices. Mutating the ALPS motif makes TPD54 cytosolic. **B**. Quantification of experiments similar to that shown in A using Superplot representation. Each dot represents one cell that is color-coded according to the biological replicate it comes from. The large circles represent the mean of each replicate. The three means were then used to calculate the average ± SEM (black bars) (57). Thirty cells from three independent experiments were analyzed for each mutant. **C**. Dynamics of GFP-TPD54 or of mutants in AH3 or in AH4 as assessed by FRAP analysis in various subcellular regions. Pure GFP was used as a control for a soluble cytosolic protein. Data show mean ± SEM from 10 to 20 FRAP assays and from two cell transfections. Scale bar: 10 µm.

Larocque et al showed that the apparent cytosolic pool of TPD54 as seen by conventional microscopy is misleading: FRAP measurements and flickering analysis of the GFP-TPD54 signal in the cytoplasm suggested association to vesicular structures (9). We performed FRAP measurements of some GFP-TPD54 constructs in selected subcellular regions, including the cytoplasm, the rounded structures and the Golgi apparatus (**Figure 5C**). Wild-type GFP-TPD54 and GFP-TPD54[V171D] recovered from these structures as well as from the cytoplasm with comparable rate constants, which were in the range of 0.2 to 0.37 s^-1^. The recovery curves, which could be well resolved, were ca 4-6 times slower than what was observed with pure GFP (1.25 s^-1^) suggesting that GFP-TPD54 was never soluble, even when present in the cytoplasmic fraction. Strikingly, when GFP-TPD54 was mutated in its ALPS motif (I151D), which led to its complete redistribution in the cytoplasm, its recovery rate was very fast (1 s^-1^), similar to that of isolated GFP (1.25 s^-1^) and much faster than wild-type GFP-TPD54, whether in the cytoplasm (0.37 s^-1^) or on visible structures (**Figure 5C**). These static and dynamics observations indicate that the ALPS motif of GFP-TPD-54 makes a major contribution to the association of the protein to various subcellular structures, including cytoplasmic structures, which were previously identified as nanovesicles.

### TPD54 lipid binding properties resemble that of other ALPS-containing proteins

Whereas the results of the limited proteolysis experiments and of NBD/Cys scanning mutagenesis suggest major rearrangements of all regions encompassing AH1 to AH4 (**Figures 2 and 4**), the cellular experiments highlight a prominent role of AH3 (ALPS) in TPD54 localization. To further investigate the role of this region in the lipid binding properties of TPD54, we combined a disruptive mutation in the ALPS motif (I151D) with NBD labelling at position S161C to follow TPD54 binding to liposomes. As a control, we prepared a second mutant harboring both the reporter mutation S161C-NBD and a disruptive mutation in AH4 (V171D).

Titration experiments with increasing concentration of sonicated PC(18:1/18:1) liposomes showed that the liposome-binding properties of TPD54[S161C-NBD] and TPD54[S161C-NBD, V171D] were similar, whereas TPD54[I151D, S161C-NBD] showed a 10-fold decrease in affinity for liposomes confirming that the ALPS motif has a predominant role in TPD54 membrane association (**Figure 6A**). In addition, the mutant bearing disruptive mutations in both AH3 and AH4 (TPD54[I151D, S161C-NBD, V171D]) showed a similar dose-response curve as the AH3 mutant, confirming the negligible role of AH4 (**Figure 6A**).

**Figure 6.**
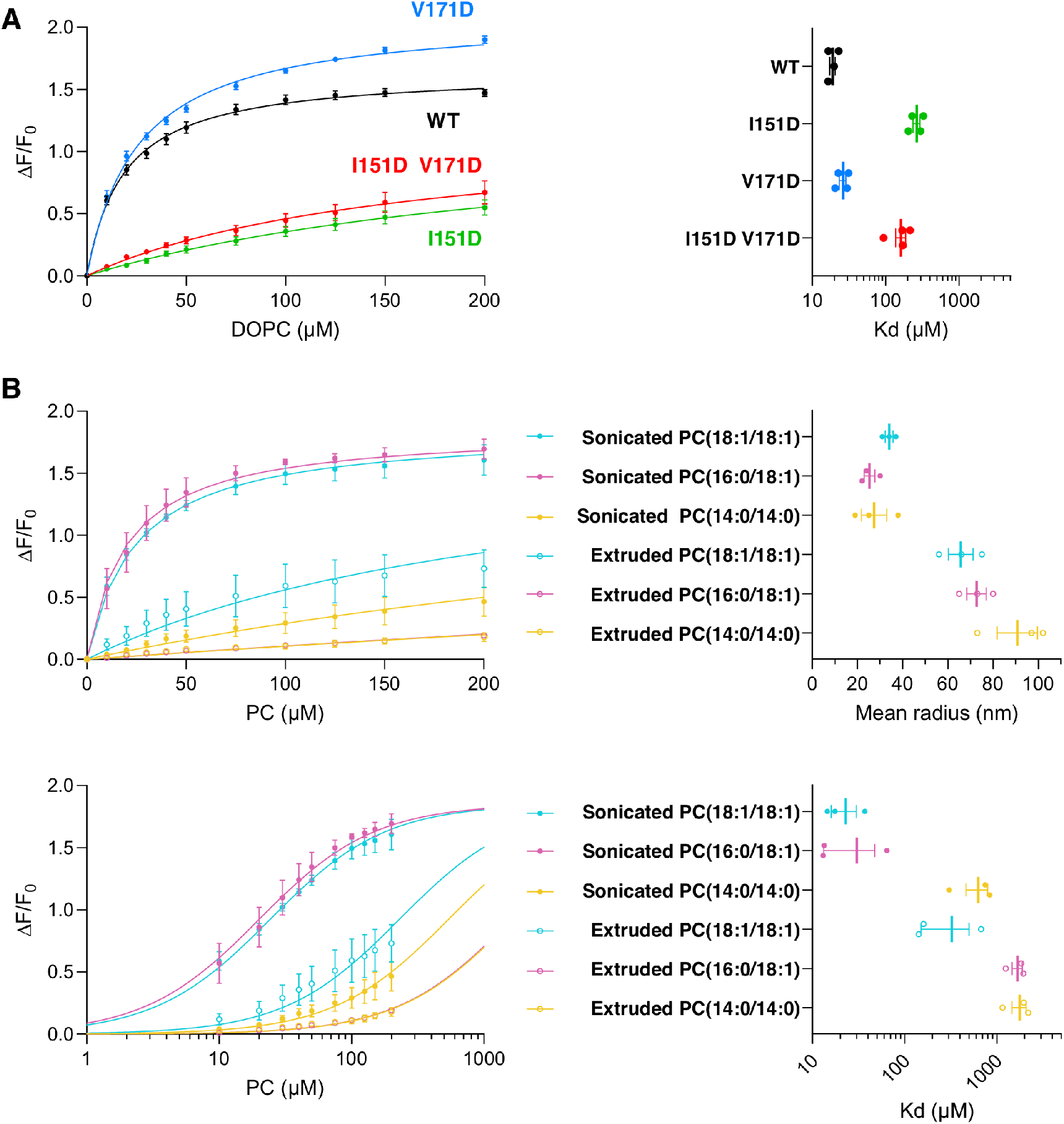
TPD54 interacts with small liposomes through its ALPS motif. **A**. Impact of disruptive mutations in the ALPS motif (AH3) or in AH4 on TPD54-S161C-NBD binding to sonicated PC(18:0:18:1) liposomes. The fluorescence at 530 nm of the protein (100 nM) was used to build liposome binding curves, which were fitted with hyperbolic functions to determine apparent K_d_. Data show mean ± SEM from 4 independent titration curves conducted with two preparations of liposomes. **B**. Dual effect of membrane curvature and lipid unsaturation on the binding of TPD54-S161C-NBD to PC liposomes. Liposomes were made of PC(18:1/18:1), PC(16:0/18:1) or PC(14:0/14:0) and were prepared by extrusion (200 nm; open symbols) or by sonication (filled symbols). The titration curves were determined from the fluorescence of TPD54-S161C-NBD (100 nM) at 530 nm and are shown on a linear (top left) or logarithm (bottom left) scale to illustrate the large differences in binding affinities. The upper right plot shows the mean size of each liposome preparation. The lower right plot shows the apparent K_d_ as deduced from hyperbolic fits. Data show mean ± SEM from 3 independent experiments conducted with different preparations of liposomes

Next, we performed titration experiments using liposomes made of either PC(18:1/18:1), PC(16:0/18:1) or PC(14:0/14:0). We observed marginal binding of TPD54[S161C-NBD] to PC(16:0/18:1) or PC(14:0/14:0) liposomes made by extrusion (Kd ≈ 2 mM; radius ≈ 70 and 90 nm, respectively) (**Figure 6B**) but more significant binding to extruded PC(18:1/18:1) liposomes (Kd ≈ 300 µM; radius ≈ 65 nm). With sonicated PC(18:1/18:1) or PC(16:0/18:1) liposomes (radius = 30 and 25 nm, respectively), binding was much stronger (Kd ≈ 20 and 25 µM; respectively), whereas barely any binding occurred with sonicated (radius 22 nm) PC(14:0/14:0) liposomes (Kd ≈ 700 µM) (**Figure 6B**). Overall, the affinity of TPD54[S161C-NBD] for PC liposomes varied by two orders of magnitude (from Kd = 20 µM to 2 mM) depending on PC unsaturation and liposome size (**Figure 6B**). This hyper sensitivity to both membrane curvature and lipid unsaturation is a biochemical feature of ALPS motifs (15, 19).

Lastly, we compared the liposome binding properties of TPD54 with other ALPS-containing constructs, namely the N-terminal region of GMAP-210 and the first ALPS motif of ArfGAP1 (**supplementary figure 4**). With PC(16:0/18:1) liposomes, GMAP-210[M1C-NBD, 1-189], ArfGAP1[192-257, A236C-NBD] and TPD54[S161C-NBD] showed binding to small liposomes (R = 39 nm) but not to larger ones (R in the 82 – 105 nm range). We also added 30% of the negatively charged lipid PS(16:0/18:1), which strongly favors the membrane binding of positively charged amphipathic helices but has minor effects on ALPS motifs (46). PS(16:0/18:1) had a slight positive effect on TPD54, the binding of which could be observed even with extruded liposomes.

These experiments indicate that TPD54 contains a functional ALPS motif, which is a major membrane-interacting region of the protein and accounts for its preference for highly curved and unsaturated membranes.

### TPD54 and GMAP-210 do not recognize the same vesicular structures

Several lines of evidence suggest that the golgin GMAP-210 recognizes endogenous vesicles through its N-terminal ALPS motif. Notably, correlative light-electron microscopy (CLEM) showed that when GMAP-210 lacked its ALPS motif, it failed to restore the Golgi ultrastructure of cells in which the expression of endogenous GMAP-210 was silenced, whereas full length GMAP-210 restored the balance between vesicles and cisternae (26). Given the involvement of an ALPS motif in the recognition of small transport vesicles by TPD54, we wondered whether GMAP-210 and TPD54 might recognize a common set of transport vesicles, which should lead to some overlap in their respective localization.

We analyzed the localization of endogenous GMAP-210 and TPD54 or of overexpressed protein constructs. Given the small size of transport vesicles and the complex organization of the Golgi apparatus, we conducted this analysis using stimulated-emission-depletion (STED) microscopy in addition to conventional confocal microscopy. When we compared STED images of endogenous TPD54 and of endogenous GMAP-210 using specific antibodies, the signals from the two proteins did not overlap, although they both stained the Golgi region (**Figure 7**). In addition, the signal of slightly overexpressed Halo-Tag-TPD54 did not coincide with that of endogenous GMAP-210. Similarly, the signal from a Halo-Tag construct that included the N-terminal ALPS motif of GMAP-210 was clearly separated from the signal of endogenous TPD-54. We concluded that GMAP-210 and TPD54 do not bind the same cellular structures despite both harboring functional ALPS motifs.

**Figure 7.**
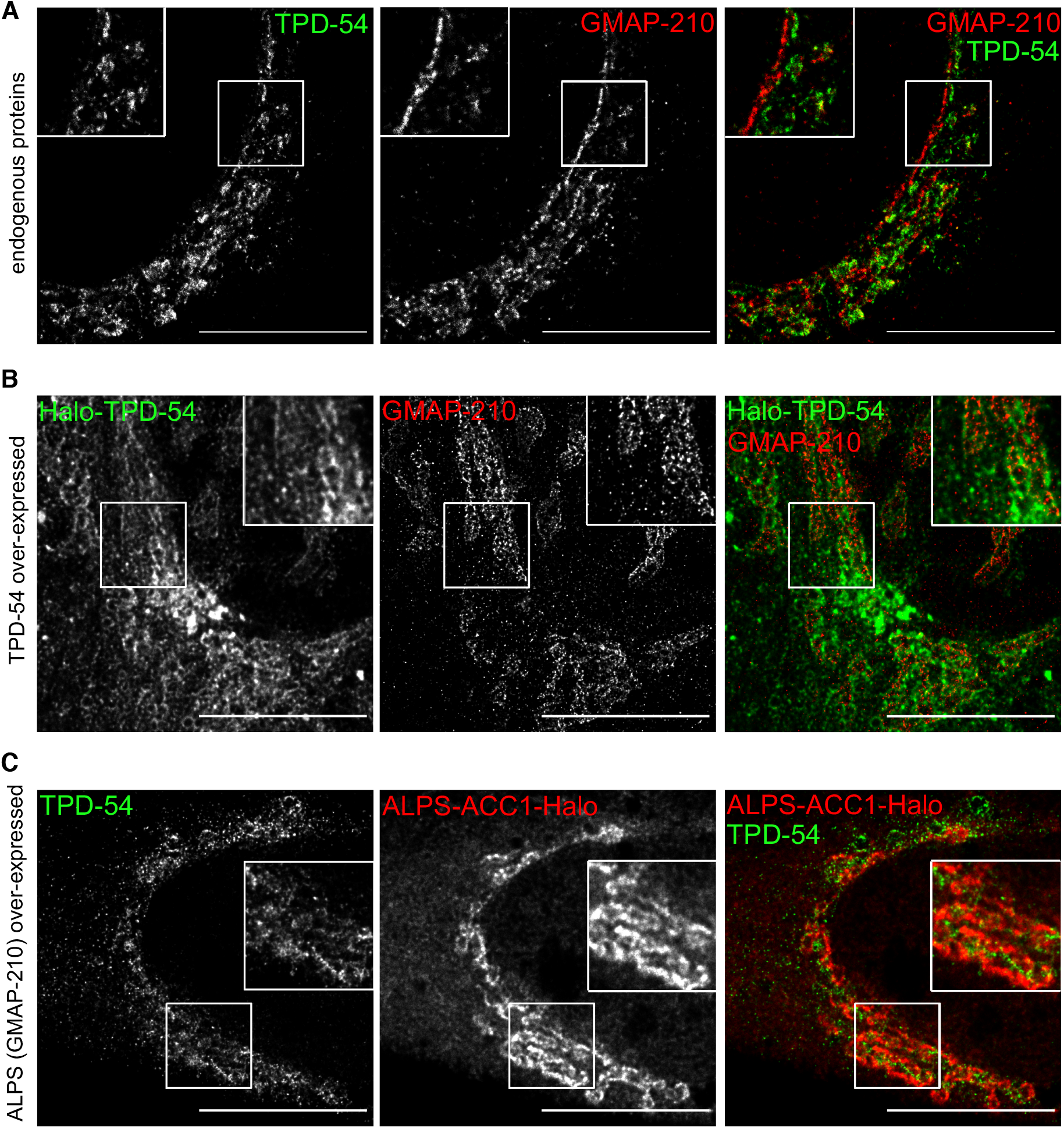
TPD54 and the golgin GMAP-210 bind different cellular structures. STED Imaging of TPD-54 as compared to GMAP-210 in RPE1 cells. **A**. Endogenous TPD-54 *vs* endogenous GMAP-210. **B**. Over-expressed Halo-TPD-54 *vs* endogenous GMAP-210. **C**. Endogenous TPD-54 *vs* a construct made of the ALPS motif of GMAP-210, an artificial coiled-coil and a Halo tag (GMAP-210-ACC1-Halo). Scale bar: 10 µm.

To further analyze the difference in localization of GMAP-210 and TPD54 and the involvement of their ALPS motifs, we prepared a chimera in which we replaced the ALPS motif of GMAP-210 with that of TPD54 in a construct corresponding to the N-terminal region of GMAP-210. This chimera was essentially soluble in marked contrast with another coiled-coil-based chimera containing the ALPS motif of GMAP-210 (14, 47), which clearly stained the Golgi apparatus (**supplementary Figure 5**). This result suggests that the ALPS motif of TPD54, although necessary for membrane binding, is not sufficient when put in a different protein context. Of note, the ALPS motif of TPD54 (18 aa) is twice as short as that of GMAP-210 (38 aa) and previous experiments showed that the entire sequence of the ALPS motif of GMAP-210 contributed to membrane binding (14).

## Discussion

The discovery that the protein TPD54 captures intracellular vesicles that are smaller than conventional transport vesicles (≈ 30 vs 60 nm in diameter) came as a surprise since it was generally assumed that the repertoire of transport vesicles in cells was complete (9). The present study, which combines an in-depth biochemical analysis of TPD54 with cellular observations, gives a molecular basis for the capture of small vesicles by TPD54.

CD spectroscopy and limited proteolysis experiments indicate that isolated TDP54 is largely disordered except for its predicted coiled-coil region. Upon binding to small liposomes, TPD54 undergoes a huge structural change, which affects the entire region downstream of the coiled-coil, i.e. from aa 85 up to amino-acid 200, and which correlates with a gain in α-helicity. In contrast, the region upstream of the coiled-coil remains unfolded and remote from the membrane. Among the regions that become folded upon membrane association, the ALPS motif of TPD54, which encompasses aa 141-158, makes a key contribution to membrane interaction. A single mutation in its hydrophobic face was sufficient to render the protein soluble in cells and lowered the avidity of purified TPD54 for small liposomes by 10-fold; a similar mutation in the subsequent amphipathic helix (AH4) had no effect.

TPD54 binds to lipid membranes in a sharp curvature- and lipid unsaturation-dependent manner. We did not detect binding of TPD54 on liposomes with a diameter > 60 nm except when using the non-physiological lipid PC(4ME-16:0/4ME-16:0), which creates large lipid-packing defects even in flat membranes (41). For liposomes made of PC(16:0/18:1), which is the most abundant PC species in cellular membranes, TPD54 shows binding only when the liposomes diameter is below 60 nm. The hypersensitivity of TPD54 to membrane curvature and lipid unsaturation contrasts with its low sensitivity to liposome charge as binding was poorly dependent on the general anionic lipid phosphatidylserine. Although we cannot exclude that TPD54 is sensitive to some specific lipids that were not tested here, the dual sensitivity of TPD54 to both membrane curvature and lipid unsaturation makes TPD54 very similar to other ALPS-motif containing proteins as regard to membrane binding properties (11, 12, 20).

Whether the ALPS motif of TPD54 has a curvature threshold different from other ALPS motifs, making it more adapted for the capture of nanovesicles with a diameter of 30 nm as compared to vesicles with a diameter of 60-80 nm, awaits further investigation. The assays used here are based on bulk measurements with liposomes prepared by sequential extrusion through polycarbonate filters or by sonication. As previously documented by electron microscopy and DLS measurements (48, 49), these methods give liposomes that display polydispersity around an average size value. To further characterize the exact curvature preference of TDP54, other methods will be necessary. These include quantitative fluorescence microscopy to measure curvature-selective binding of protein on immobilized single liposomes (50, 51) or on tubes pulled from giant unilamellar vesicles (22, 52).

The fact that structuration of TPD54 is not restricted to its ALPS motif and that the subcellular localization of TPD54 differs from that of the golgin GMAP-210, which also recognizes transport vesicles through an ALPS motif, suggest that other determinants cooperate with the ALPS motif to make TPD54 and the golgin GMAP-210 specific to different populations of vesicles. In the case of GMAP-210, both the ALPS motif and the very first N-terminal residues are involved in the capture of different vesicles (53). For TPD54, a recent study based on the localization of various GFP constructs and on the effect of FKPB-TPD54 constructs on mitochondria aggregation suggests that region 83-126 and a set of basic residues (K154, R159, K175, and K177) are involved in intracellular nanovesicle recognition, although the underlying mechanism is not known (54). Determining how these various regions and the ALPS motif of TPD54 cooperate to selectively capture INVs through protein-protein and protein-membrane interactions will require numerous investigations because their disordered nature makes structural investigations difficult. α-synuclein and CCTα provide nice examples of such efforts where extensive mutagenesis coupled to spectroscopic methods (e.g. fluorescence and EPR) enable to establish the geometry of these amphitropic proteins at the membrane surface (55, 56). We considered unlikely that all four predicted amphipathic helices of TPD54 (from AH1 to AH4) directly contribute to lipid membrane interactions as this would require strong distortion in the regions separating them to allow their hydrophobic faces to point simultaneously toward the membrane. Nevertheless, the identification of a functional ALPS motif in TPD54 gives a first hint for the mechanism of selective capture of very small transport vesicles by this protein.

## Experimental procedures

### Molecular biology

The TPD54 sequence was amplified by from Hela cell RNA extracts by RT PCR. The TPD54 sequence was subcloned into a pGEX-2T vector at the BamH1 and EcoR1 sites, i.e. following the GST and the Thrombin cleavage site. Mutations were introduced with the QuickChange Lightning kit (Agilent).

### Protein expression and purification

We expressed TPD54 as a GST fusion in E Coli BL21. Bacteria were grown in LB supplemented with 100µg/ml ampicillin at 37°C. Protein expression was induced by the addition of 1 mM IPTG at OD = 0.8 for 1h30’ at 37°C. Bacteria were lysed in a cell disruptor at 1600 psi in Tris 50 mM pH 7.4, NaCl 150 mM, DTT 2 mM. After addition of 0.1 mg/ml DNAse and 5 mM MgCl_2_, the soluble fraction was collected by centrifugation at 40 000 rpm for 30 min in a Ti45 rotor (Beckman). The fusion protein was purified from the soluble fraction using glutathione-Sepharose 4B beads at a vol/vol ratio of 4 ml beads for 100 ml supernatant. TPD54 was released from the beads by thrombin cleavage (12U/ml in the presence 50 µM CaCl_2_) of the linker between GST and TPD54, which leaves the sequence “GS” at the N-terminus of the protein. TPD54 was further purified by gel-filtration on a Sephadex S300 HR column in 20 mM Tris pH 7.4, 150 mM NaCl, 10% vol/vol Glycerol and 1 mM DTT. Aliquots were flash-frozen in liquid nitrogen and stored at -80°C. Purification of ArfGAP1 ALPS1 motif [192-257, A236C-NBD] has been described elsewhere (15, 21). GMAP-210 [M1C-NBD, 1-189] was prepared from our previous construct GMAP-210 [M1C-NBD, 1-375] and purified and labelled with the same protocol (13).

### Protein labelling with fluorescent probes

Cysteine mutants of TPD54 were labelled with NBD using a 10-fold excess of IANBD Amide (N,N’-Dimethyl-N-(Iodoacetyl)-N’-(7-Nitrobenz-2-Oxa-1,3-Diazol-4-yl)Ethylenediamine) in 50 mM Tris pH 7.4, 150 mM NaCl. The time and temperature of labelling was optimized to limit the labelling of the sole endogenous cysteine (C74), which was found to have limited reactivity due to its intra-coiled-coil localization. Excess probe was blocked with L-cysteine and removed by size exclusion chromatography. The concentration of labelled protein was determined by absorbance at 500 nm.

### Liposome preparation

Lipids were mixed as chloroform solutions and dried in a rotary evaporator. The lipid film was resuspended in HK buffer (50 mM HEPES pH 7.2, 120 mM K acetate) and the suspension was frozen and thawed 5 times to promote the formation of unilamellar liposomes followed by sequential extrusion through polycarbonate filters of decreasing pore size (200, 100, 50 and 30 nm). To further decrease the liposome size, the liposome suspension was sonicated using a probe sonicator. The average liposome size and polydispersity were determined by dynamic light scattering using a Cordouan Vasco Kin Particle Size Analyzer. As small liposomes have tendency to fuse, they were used within the same day of preparation.

### Liposome flotation assay

Liposomes and TPD54 were incubated for 5 min in 50 mM HEPES pH 7.2, 120 mM K acetate, 1 mM MgCl_2_ (HKM buffer) at room temperature in a Beckman polycarbonate centrifuge tube. The mixture (75 µl) was mixed with HK buffer supplemented with 60% sucrose (100 µl) and then overlaid with an intermediate sucrose solution made of HK buffer with 24% sucrose (100 µl) and a layer (25 µl) of HKM buffer. After centrifugation at 100 000 rpm in a fixed-angle TLA100 rotor for 1 hr, three fractions (bottom, medium, and top) of 175, 75 and 50 µl were collected and then analyzed by SDS-PAGE using Sypro-Orange staining.

### Limited proteolysis

Proteins (2µM) were mixed in HKM buffer with or without sonicated PC(18:1/18:1) liposomes at a protein/lipid mol:mol ratio of 1/1000 to get maximal membrane binding of TPD54. Trypsin or subtilisin (1 µg/ml) was added at time zero under agitation. At the indicated times, aliquots were withdrawn and the reaction was stopped by adding 0.5 µM PMSF. The reaction was analyzed by SDS Page with Sypro-Orange staining and by mass spectrometry.

### Mass spectrometry analysis - µHPLC-Q-exactive plus

The sample was analyzed by liquid chromatography coupled to a mass spectrometer equipped with a heated electrospray ionization (HESI) probe. HPLC was performed using a Dionex U3000 RSCL Instrument. The injection volume was fixed at 5 µl (Ultimate 3000, Thermo Fisher Scientific). Protein analysis was performed on a 2.1mm i.d. x 150mm, 5µm, Syncronis C18 column at a flow rate of 250 µl/min. The elution program was based on water (solvent A) and acetonitrile (solvent B) both containing 0.1% formic acid (v/v): 0 min 5% B, 18 min 80% B, 19 min 90% B. The Q-exactive plus spectrometer completely controlled by the Xcalibur software was operating in electrospray positive mode. Typical ESI conditions were as follows: electrospray voltage 4 kV; capillary temperature 320°C, probe temperature 325°C, sheath gas flow 40U and auxiliary gas 10U. The MS scan was acquired in the 400 to 1900 m/z range with the resolution set to 140 000. Data analysis was performed with Biopharma Finder 3.2 and intact protein spectra was automatically deconvoluted with Default-Auto Xtract.

### CD spectroscopy

The protein was dialyzed against Potassium Phosphate Fluoride buffer (10mM KH_2_PO_4_, 100 mM KF, 0.1 mM DTT) and then supplemented with sonicated DOPC liposomes at increasing concentration in a quartz cell with an optical path length of 0.05 cm. CD spectra were measured at room temperature in a Jasco J-815 CD spectrometer at a scan speed of 50nm/min from 195 to 260nm. For each condition, 5 spectra were accumulated and corrected for the blank, which was recorded under the same conditions in the absence of protein. Estimation of the percentage of α-helix was performed using the CD analysis and plotting tool CAPITO (https://data.nmr.uni-jena.de/capito/index.php).

### Fluorescence experiments

All fluorescence experiments were performed in a Jasco FP 8300 fluorimeter using a cylindrical cuvette (volume 600 µl) equipped with a stir bar and thermostated at 37°C. For NBD fluorescence, excitation was set at 500 nm (bandwidth 1 nm) and emission was recorded from 520 to 650 nm (bandwidth 10 nm). For tryptophan fluorescence, excitation was set at 295 nm (bandwidth 2.5 nm) and emission was recorded from 305 to 450 nm (bandwidth 5 nm). Fluorescence energy transfer between TPD54 tryptophans and DPH-PC was measured by exciting the tryptophans at 280 nm (bandwidth 1 nm) and following the emission of DPH-PC at 460 nm (bandwidth 5 nm) over time. The cuvette initially contained HKM buffer (Hepes 50 mM, pH 7.2, K acetate 120 mM, MgCl_2_ 1 mM, DTT 1 mM). For NBD fluorescence, 0.1 µM NBD-labeled TPD54 was added followed by liposomes of defined size and composition. For tryptophan fluorescence, 0.1 µM unlabeled TPD54 was added followed by liposomes of defined size and composition. Emission spectra were recorded after each addition and were corrected for the corresponding blank, i.e. a similar buffer/liposome mixture in the absence of protein. The proximity of tryptophans to liposomes was measured by FRET using DPH-PC-containing liposomes. Increasing amounts of unlabeled TPD54 were added to the liposome suspension.

### Cell culture, protein expression and antibodies

hTERT-RPE1 cells from ATCC were cultured in DMEM/F12 medium with glutaMAX (Gibco) containing 10% serum, 1% antibiotics (Zell Shield, Minerva Biolabs) and were incubated in a 5% CO2 humidified atmosphere at 37°C. Cells were seeded on µDish or µSlide (Ibidi). Plasmids were transfected using Lipofectamine 3000 reagent according to the instruction’s guidelines (ThermoFisher Scientific). After 6 hours, the cells were fixed with PFA 4% for 20 min at room temperature. Golgi staining was performed using sheep polyclonal antibody against TGN46 (AHP500G, Bio-Rad).

Live cell observations for FRAP experiments were done using a confocal microscope (Zeiss LSM-780). Images were analyzed using Fiji J software.

To determine the fraction of GFP-TPD54 and mutants at the Golgi, we outlined the Golgi area from the TGN46 signal carried out in the red channel. Then, we applied the corresponding mask to the green channel to define a region of interest (ROI) at the Golgi and a ROI of the same area in the cytosol. Once the background was subtracted, we determined the average fluorescence in each ROI and calculated the Golgi/cytosol ratio. All images were acquired with a confocal microscope and were analyzed using Volocity software.

For STED experiments, TPD-54 and ALPS(GMAP-210)-ACC1 were cloned in the halo-tag vectors HTN and HTC, respectively. After transfection, these constructs were revealed using a Janelia Fluor 549 ligand (promega). Endogeneous proteins were labeled with Rb polyclonal antibody anti-TPD52L2 (proteintech), Rb polyclonal antibody anti-GMAP-210 (proteintech) or Ms monoclonal anti-Trip230 human (Invitrogen). Images were acquired on a Leica confocal TCS SP8 STED microscope. Deconvolution was performed using Huygens Professional software.

## Abbreviations

AH: (amphipathic helix)
ALPS: (Amphipathic Lipid Packing Sensor)
CD: (circular dichroism)
DPH: (diphenylhexatriene)
ER: (endoplasmic reticulum)
GA: (Golgi apparatus)
NBD: (Nitrobenzoxadiazole)
PC: (phosphatidylcholine)
PC(18:1/18:1): (dioleoyl phosphatidylcholine)
PC(16:0/18:1): (1-palmitoyl 2-oleoyl phosphatidylcholine)
PC(14:0/14:0): (dimyristoyl phosphatidylcholine)
PC(4ME 16:0/4ME 16:0): (diphytanoyl phosphatidylcholine)
PS: (phosphatidylserine)
TGN: (trans Golgi network)
TPD54: (Tumor Protein D54).

## Acknowledgements

We thank S Miserey (institute Curie), G Lenoir (Université Paris Saclay) and all lab members for support and helpful discussions.

## Author contributions

A.R. and M.M. conceptualization, investigation, methodology, data curation, formal analysis, visualization, writing—review and editing. S.A., A.S.G and D.D methodology and data curation. A.P. methodology, writing—review and editing. B.A. conceptualization, supervision, writing— original draft, writing—review and editing, and funding acquisition.

## Funding and additional information

This work was supported by the Agence Nationale de la Recherche (ANR Grant #n° 195624, Vesicle Filtering).

## Data availability

All data are contained within the article and the supporting information.

## Supporting information

Supplemental Figures S1–S4

## References

1. Bonifacino, J. S., and Glick, B. S. (2004) The mechanisms of vesicle budding and fusion. Cell. 116, 153–166

2. Hurley, J. H., Boura, E., Carlson, L.-A., and Róźycki, B. (2010) Membrane Budding. Cell. 143, 875–887

3. Dell’Angelica, E. C., and Bonifacino, J. S. (2019) Coatopathies: Genetic Disorders of Protein Coats. Annu. Rev. Cell Dev. Biol. 35, 131–168

4. Lee, M. C. S., Miller, E. A., Goldberg, J., Orci, L., and Schekman, R. (2004) Bi-directional protein transport between the ER and Golgi. Annu. Rev. Cell Dev. Biol. 20, 87–123

5. Barlowe, C. (1994) COPII: A membrane coat formed by Sec proteins that drive vesicle budding from the endoplasmic reticulum. Cell. 77, 895–907

6. Malhotra, V., Serafini, T. A., Orci, L., Shepherd, J. C., and Rothman, J. E. (1989) Purification of a novel class of coated vesicles mediating biosynthetic protein transport through the Golgi stack. Cell. 58, 329–336

7. McCaughey, J., and Stephens, D. J. (2019) ER-to-Golgi Transport: A Sizeable Problem. Trends in Cell Biology. 29, 940–953

8. Hutchings, J., and Zanetti, G. (2019) Coat flexibility in the secretory pathway: a role in transport of bulky cargoes. Current Opinion in Cell Biology. 59, 104–111

9. Larocque, G., La-Borde, P. J., Clarke, N. I., Carter, N. J., and Royle, S. J. (2019) Tumor protein D54 defines a new class of intracellular transport vesicles. The Journal of Cell Biology. 10.1083/jcb.201812044

10. Boutros, R., Fanayan, S., Shehata, M., and Byrne, J. A. (2004) The tumor protein D52 family: many pieces, many puzzles. Biochem. Biophys. Res. Commun. 325, 1115–1121

11. Giménez-Andrés, M., Čopič, A., and Antonny, B. (2018) The Many Faces of Amphipathic Helices. Biomolecules. 8, 45

12. Drin, G., and Antonny, B. (2010) Amphipathic helices and membrane curvature. FEBS Letters. 584, 1840–1847

13. Drin, G., Casella, J.-F., Gautier, R., Boehmer, T., Schwartz, T. U., and Antonny, B. (2007) A general amphipathic alpha-helical motif for sensing membrane curvature. Nat Struct Mol Biol. 14, 138–146

14. Magdeleine, M., Gautier, R., Gounon, P., Barelli, H., Vanni, S., and Antonny, B. (2016) A filter at the entrance of the Golgi that selects vesicles according to size and bulk lipid composition. eLife. 5, 292

15. Bigay, J., Casella, J.-F., Drin, G., Mesmin, B., and Antonny, B. (2005) ArfGAP1 responds to membrane curvature through the folding of a lipid packing sensor motif. The EMBO Journal. 24, 2244–2253

16. Vanni, S., Vamparys, L., Gautier, R., Drin, G., Etchebest, C., Fuchs, P. F. J., and Antonny, B. (2013) Amphipathic Lipid Packing Sensor Motifs: Probing Bilayer Defects with Hydrophobic Residues. Biophysj. 104, 575–584

17. van Hilten, N., Stroh, K. S., and Risselada, H. J. (2020) Membrane Thinning Induces Sorting of Lipids and the Amphipathic Lipid Packing Sensor (ALPS) Protein Motif. Front Physiol. 11, 250

18. Vamparys, L., Gautier, R., Vanni, S., Bennett, W. F. D., Tieleman, D. P., Antonny, B., Etchebest, C., and Fuchs, P. F. J. (2013) Conical Lipids in Flat Bilayers Induce Packing Defects Similar to that Induced by Positive Curvature. Biophysj. 104, 585–593

19. Vanni, S., Hirose, H., Barelli, H., Antonny, B., and Gautier, R. (2014) A sub-nanometre view of how membrane curvature and composition modulate lipid packing and protein recruitment. Nature Communications. 5, 4916

20. Antonny, B. (2011) Mechanisms of Membrane Curvature Sensing. Annu. Rev. Biochem. 80, 101–123

21. Mesmin, B., Drin, G., Levi, S., Rawet, M., Cassel, D., Bigay, J., and Antonny, B. (2007) Two Lipid-Packing Sensor Motifs Contribute to the Sensitivity of ArfGAP1 to Membrane Curvature †. Biochemistry. 46, 1779–1790

22. Ambroggio, E. E., Sorre, B., Bassereau, P., Goud, B., Manneville, J.-B., and Antonny, B. (2010) ArfGAP1 generates an Arf1 gradient on continuous lipid membranes displaying flat and curved regions. The EMBO Journal. 29, 292–303

23. Zendeh-boodi, Z., Yamamoto, T., Sakane, H., and Tanaka, K. (2013) Identification of a second amphipathic lipid-packing sensor-like motif that contributes to Gcs1p function in the early endosome-to-TGN pathway. J. Biochem. 153, 573–587

24. Xu, P., Baldridge, R. D., Chi, R. J., Burd, C. G., and Graham, T. R. (2013) Phosphatidylserine flipping enhances membrane curvature and negative charge required for vesicular transport. The Journal of Cell Biology. 202, 875–886

25. Drin, G., Morello, V., Casella, J.-F., Gounon, P., and Antonny, B. (2008) Asymmetric tethering of flat and curved lipid membranes by a golgin. Science. 320, 670–673

26. Sato, K., Roboti, P., Mironov, A. A., and Lowe, M. (2015) Coupling of vesicle tethering and Rab binding is required for in vivo functionality of the golgin GMAP-210. Mol. Biol. Cell. 26, 537–553

27. Cabrera, M., Langemeyer, L., Mari, M., Rethmeier, R., Orban, I., Perz, A., Bröcker, C., Griffith, J., Klose, D., Steinhoff, H.-J., Reggiori, F., Engelbrecht-Vandré, S., and Ungermann, C. (2010) Phosphorylation of a membrane curvature-sensing motif switches function of the HOPS subunit Vps41 in membrane tethering. The Journal of Cell Biology. 191, 845–859

28. Ho, R., and Stroupe, C. (2016) The HOPS/Class C Vps Complex Tethers High-Curvature Membranes via a Direct Protein-Membrane Interaction. Traffic. 17, 1078–1090

29. Krabben, L., Fassio, A., Bhatia, V. K., Pechstein, A., Onofri, F., Fadda, M., Messa, M., Rao, Y., Shupliakov, O., Stamou, D., Benfenati, F., and Haucke, V. (2011) Synapsin I senses membrane curvature by an amphipathic lipid packing sensor motif. J. Neurosci. 31, 18149–18154

30. Moser von Filseck, J., Vanni, S., Mesmin, B., Antonny, B., and Drin, G. (2015) A phosphatidylinositol-4-phosphate powered exchange mechanism to create a lipid gradient between membranes. Nature Communications. 6, 6671

31. Monje-Galvan, V., and Klauda, J. B. (2018) Preferred Binding Mechanism of Osh4’s Amphipathic Lipid-Packing Sensor Motif, Insights from Molecular Dynamics. J Phys Chem B. 122, 9713–9723

32. Brier, L. W., Ge, L., Stjepanovic, G., Thelen, A. M., Hurley, J. H., and Schekman, R. (2019) Regulation of LC3 lipidation by the autophagy-specific class III phosphatidylinositol-3 kinase complex. Mol. Biol. Cell. 30, 1098–1107

33. Fan, W., Nassiri, A., and Zhong, Q. (2011) Autophagosome targeting and membrane curvature sensing by Barkor/Atg14(L). Proc. Natl. Acad. Sci. U.S.A. 108, 7769–7774

34. Ohashi, Y., Tremel, S., Masson, G. R., McGinney, L., Boulanger, J., Rostislavleva, K., Johnson, C. M., Niewczas, I., Clark, J., and Williams, R. L. (2020) Membrane characteristics tune activities of endosomal and autophagic human VPS34 complexes. eLife. 10.7554/eLife.58281

35. Mesmin, B., Bigay, J., Polidori, J., Jamecna, D., Lacas-Gervais, S., and Antonny, B. (2017) Sterol transfer, PI4P consumption, and control of membrane lipid order by endogenous OSBP. The EMBO Journal. 36, 3156–3174

36. Nordeen, S. A., Turman, D. L., and Schwartz, T. U. (2020) Yeast Nup84-Nup133 complex structure details flexibility and reveals conservation of the membrane anchoring ALPS motif. Nature Communications. 11, 6060

37. Sathasivam, P., Bailey, A. M., Crossley, M., and Byrne, J. A. (2001) The role of the coiled-coil motif in interactions mediated by TPD52. Biochem. Biophys. Res. Commun. 288, 56–61

38. Tunyasuvunakool, K., Adler, J., Wu, Z., Green, T., Zielinski, M., Žídek, A., Bridgland, A., Cowie, A., Meyer, C., Laydon, A., Velankar, S., Kleywegt, G. J., Bateman, A., Evans, R., Pritzel, A., Figurnov, M., Ronneberger, O., Bates, R., Kohl, S. A. A., Potapenko, A., Ballard, A. J., Romera-Paredes, B., Nikolov, S., Jain, R., Clancy, E., Reiman, D., Petersen, S., Senior, A. W., Kavukcuoglu, K., Birney, E., Kohli, P., Jumper, J., and Hassabis, D. (2021) Highly accurate protein structure prediction for the human proteome. Nature. 10.1038/s41586-021-03828-1

39. Gautier, R., Douguet, D., Antonny, B., and Drin, G. (2008) HELIQUEST: a web server to screen sequences with specific alpha-helical properties. Bioinformatics. 24, 2101–2102

40. Wong, M., and Munro, S. (2014) Membrane trafficking. The specificity of vesicle traffic to the Golgi is encoded in the golgin coiled-coil proteins. Science. 346, 1256898–1256898

41. Garten, M., Prévost, C., Cadart, C., Gautier, R., Bousset, L., Melki, R., Bassereau, P., and Vanni, S. (2015) Methyl-branched lipids promote the membrane adsorption of α-synuclein by enhancing shallow lipid-packing defects. Phys Chem Chem Phys. 17, 15589–15597

42. Rath, A., Glibowicka, M., Nadeau, V. G., Chen, G., and Deber, C. M. (2009) Detergent binding explains anomalous SDS-PAGE migration of membrane proteins. Proc. Natl. Acad. Sci. U.S.A. 106, 1760–1765

43. Wiedemann, C., Bellstedt, P., and Görlach, M. (2013) CAPITO--a web server-based analysis and plotting tool for circular dichroism data. Bioinformatics. 29, 1750–1757

44. Antonny, B., Béraud-Dufour, S., Chardin, P., and Chabre, M. (1997) N-terminal hydrophobic residues of the G-protein ADP-ribosylation factor-1 insert into membrane phospholipids upon GDP to GTP exchange. Biochemistry. 36, 4675–4684

45. Jamecna, D., Polidori, J., Mesmin, B., Dezi, M., Levy, D., Bigay, J., and Antonny, B. (2019) An Intrinsically Disordered Region in OSBP Acts as an Entropic Barrier to Control Protein Dynamics and Orientation at Membrane Contact Sites. Developmental Cell. 10.c1016/j.devcel.2019.02.021

46. Pranke, I. M., Morello, V., Bigay, J., Gibson, K., Verbavatz, J. M., Antonny, B., and Jackson, C. L. (2011) -Synuclein and ALPS motifs are membrane curvature sensors whose contrasting chemistry mediates selective vesicle binding. The Journal of Cell Biology. 194, 89–103

47. Horchani, H., de Saint-Jean, M., Barelli, H., and Antonny, B. (2014) Interaction of the Spo20 Membrane-Sensor Motif with Phosphatidic Acid and Other Anionic Lipids, and Influence of the Membrane Environment. PLoS ONE. 9, e113484–22

48. Mayer, L. D., Hope, M. J., and Cullis, P. R. (1986) Vesicles of variable sizes produced by a rapid extrusion procedure. Biochim. Biophys. Acta. 858, 161–168

49. Olson, F., Hunt, C. A., Szoka, F. C., Vail, W. J., and Papahadjopoulos, D. (1979) Preparation of liposomes of defined size distribution by extrusion through polycarbonate membranes. Biochim. Biophys. Acta. 557, 9–23

50. Hatzakis, N. S., Bhatia, V. K., Larsen, J., Madsen, K. L., Bolinger, P.-Y., Kunding, A. H., Castillo, J., Gether, U., Hedegård, P., and Stamou, D. (2009) How curved membranes recruit amphipathic helices and protein anchoring motifs. Nature Chemical Biology. 5, 835–841

51. Jensen, M. B., Bhatia, V. K., Jao, C. C., Rasmussen, J. E., Pedersen, S. L., Jensen, K. J., Langen, R., and Stamou, D. (2011) Membrane curvature sensing by amphipathic helices: a single liposome study using α-synuclein and annexin B12. J. Biol. Chem. 286, 42603–42614

52. Tian, A., and Baumgart, T. (2009) Sorting of lipids and proteins in membrane curvature gradients. Biophys. J. 96, 2676–2688

53. Wong, M., Gillingham, A. K., and Munro, S. (2017) The golgin coiled-coil proteins capture different types of transport carriers via distinct N-terminal motifs. BMC Biol. 15, 3

54. Larocque, G., Moore, D. J., Sittewelle, M., Kuey, C., Hetmanski, J. H. R., La-Borde, P. J., Wilson, B. J., Clarke, N. I., Caswell, P. T., and Royle, S. J. (2021) Intracellular nanovesicles mediate α5β1 integrin trafficking during cell migration. The Journal of Cell Biology. 10.1083/jcb.202009028

55. Jao, C. C., Hegde, B. G., Chen, J., Haworth, I. S., and Langen, R. (2008) Structure of membrane-bound alpha-synuclein from site-directed spin labeling and computational refinement. Proc. Natl. Acad. Sci. U.S.A. 105, 19666–19671

56. Knowles, D. G., Lee, J., Taneva, S. G., and Cornell, R. B. (2019) Remodeling of the interdomain allosteric linker upon membrane binding of CCTÎ± pulls its active site close to the membrane surface. Journal of Biological Chemistry. 294, 15531–15543

57. Goedhart, J. (2021) SuperPlotsOfData-a web app for the transparent display and quantitative comparison of continuous data from different conditions. Mol. Biol. Cell. 32, 470–474

