## supplementary information for "Tumor protein D54 binds intracellular nanovesicles *via* an amphipathic lipid packing sensor (ALPS) motif"

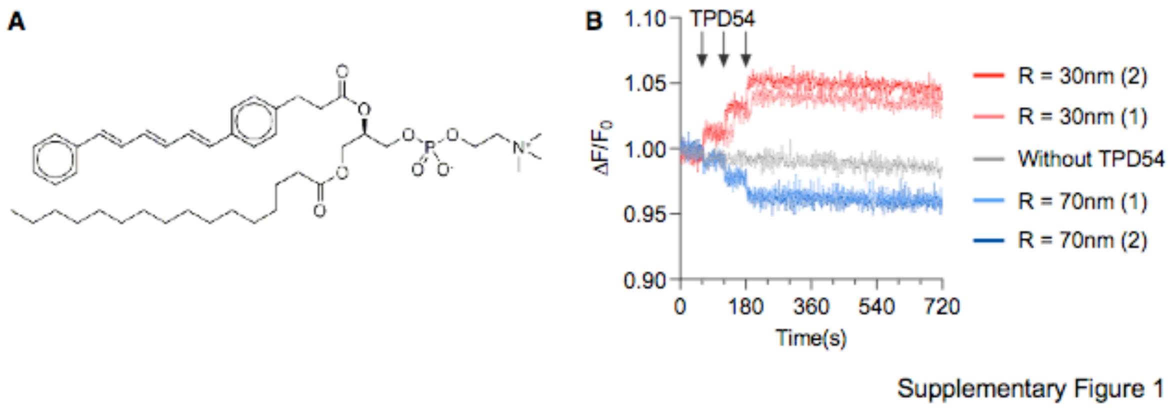


**Supplementary Figure 1. FRET assay between TPD54 tryptophan residues and DPH-PC. A.** Structure of DPH-PC. **B.** The fluorescence cuvette initially contained PC liposomes (200 µM total lipids; PC(18:1/18:1) 95 mol%, DPH-PC 5 mol%) in HKM buffer. Fluorescence at 460 nm (bandwidth 5 nm) upon excitation at 280 nm (bandwidth 1 nm) was continuously recorded upon the sequential addition of TPD54 (3 x 250 nM; vertical arrows). The red and blue traces show two independent experiments performed with sonicated liposomes (radius 30 nm) and extruded (200 nm) liposomes (radius 70 nm), respectively. The grey trace shows the evolution of the fluorescence signal without any protein addition. Note that each protein addition dilutes the sample by 1.3 %. This dilution factor was not corrected and explained the drop observed upon extruded liposome addition.

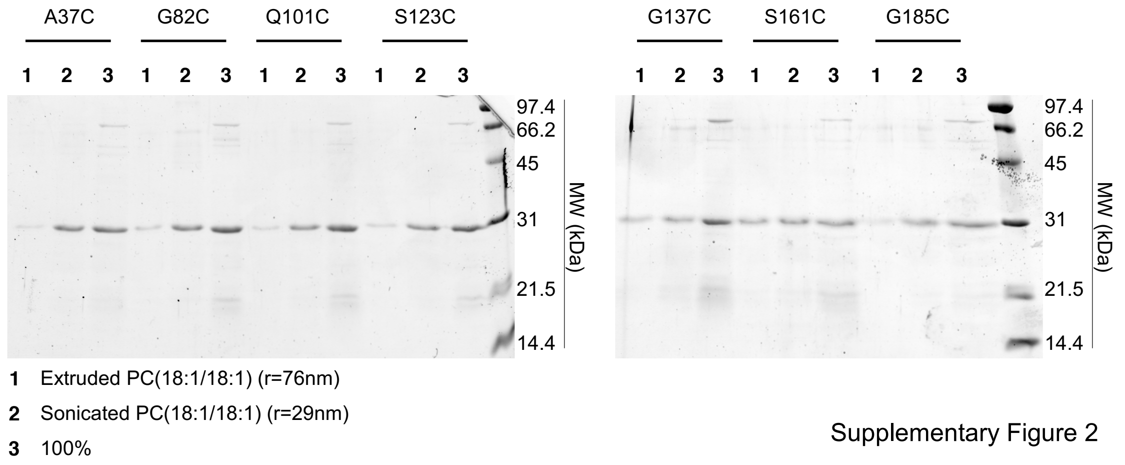


**Supplementary Figure 2. Flotation experiments of the various Cys mutants of TPD54.** After incubation of purified wild-type TPD54 or the indicated Cys mutants with extruded (200 nm) or sonicated DOPC liposomes, the liposomes were recovered by flotation on sucrose cushions and analyzed by SDS-PAGE using Sypro orange staining. The hydrodynamic radius of the extruded (200 nm) and sonicated DOPC liposomes was 76 nm and 29 nm, respectively.

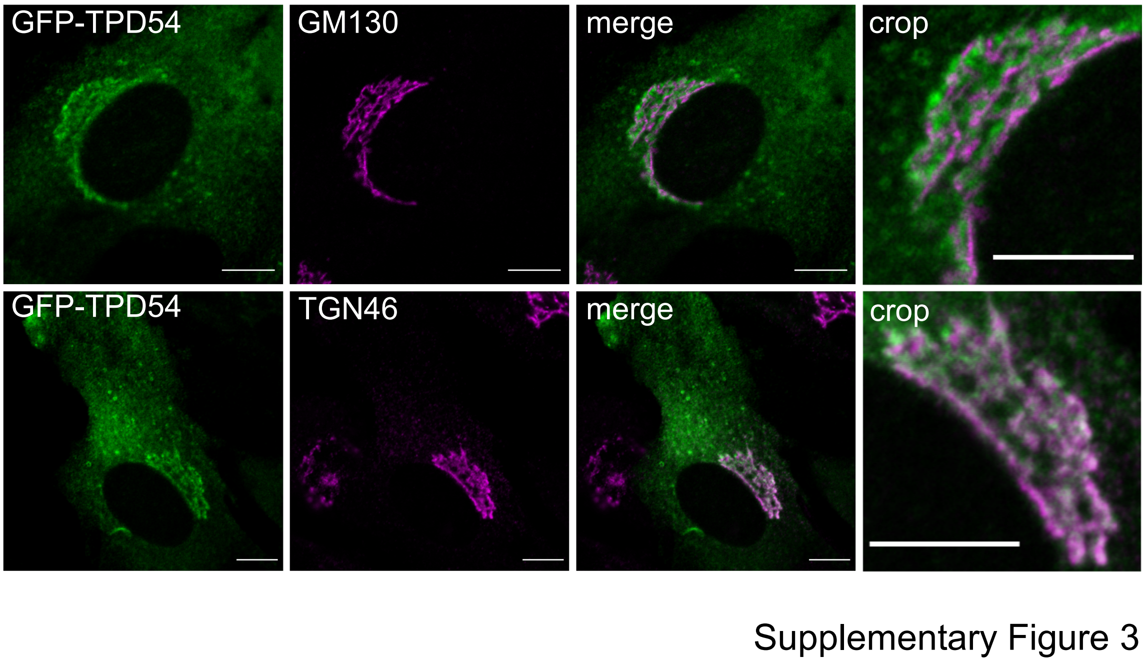


**Supplementary Figure 3. Subcellular localization of TPD54.** Confocal images of RPE1 cells expressing GFP-TPD54 and processed for immunofluorescence using antibodies against *cis* (GM130) and *trans* (TGN46) Golgi markers. Scale bar = 10 µm.

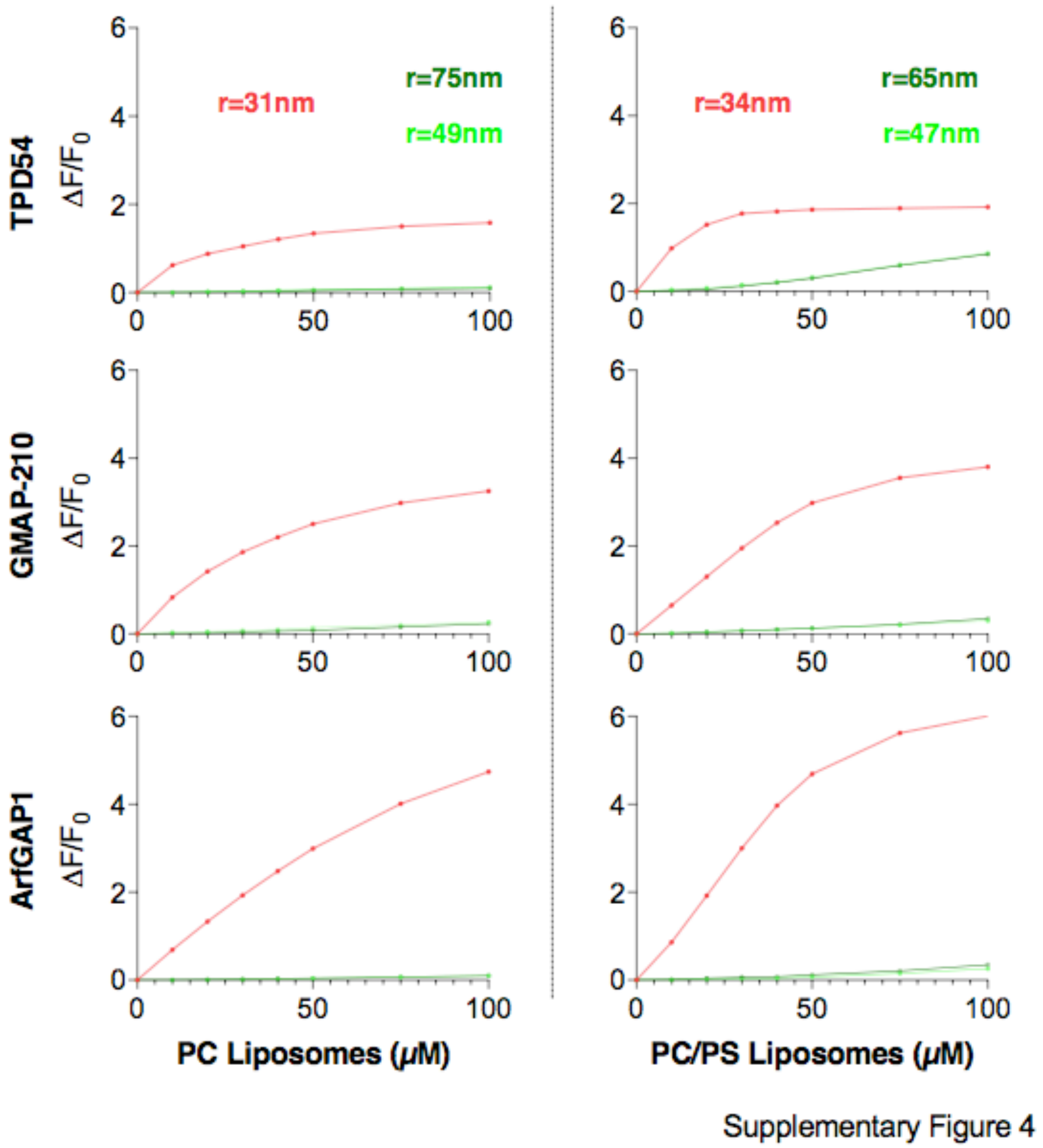


**Supplementary Figure 4. Comparison of the liposome binding properties of TPD54, the N-terminal region of GMAP-210 and the ALPS1 motif of ArfGAP1. The fluorescente cuvette initially contained 100 nM** TPD54-S161C-NBD, GMAP-210 [1-189 M1C-NBD] or ArfGAP1 [192-257, A236C-NBD]. The NBD fluorescence at 530 nm was measured with increasing amounts of extruded (green curves) or sonicated (red curves) liposomes. The experiments on the left were performed with PC(16:0/18:1) liposomes. The experiments on the right were performed with PC(16:0/18:1)/PS(16:0/18:1) (70/30) liposomes.

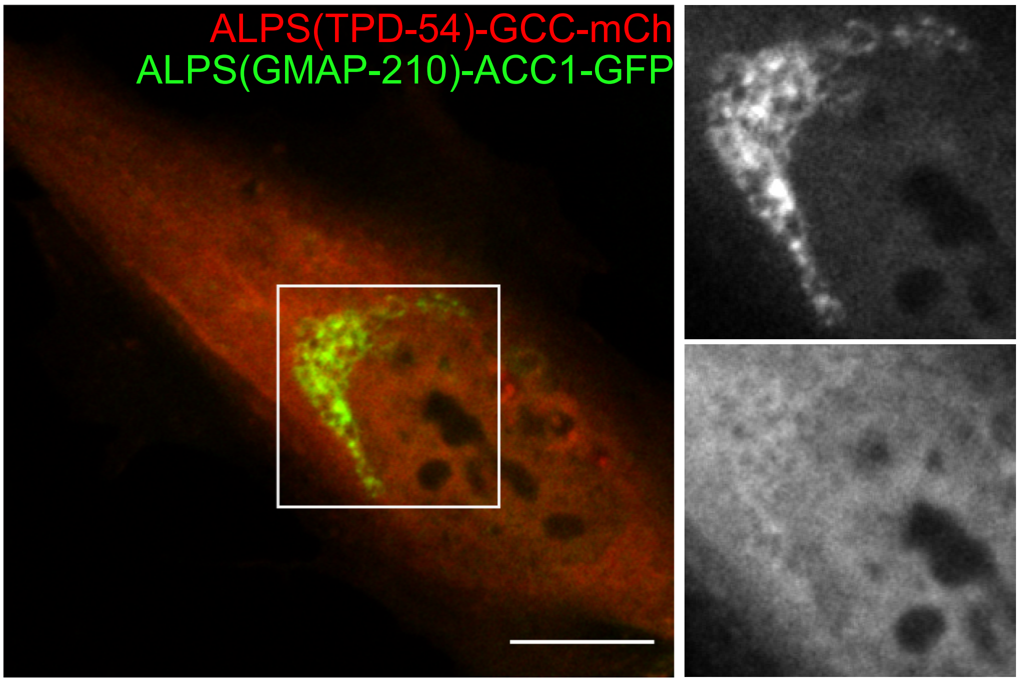


Supplementary Figure 5

**Supplementary Figure 5. A chimera construct made of the ALPS motif of TPD54 and the N-terminal region of GMAP-210 is cytosolic.** Comparison of the subcellular distribution of two fluorescent chimera in RPE1 cells. ALPS(TP54)-GCC-mCherry consists of the ALPS motif of TPD54 (aa 141-158) part of the coiled-coil region of GMAP (aa 39-375) and mCherry as a fluorescent reporter. ALPS(GMAP-210)-ACC1-GFP consists of the ALPS motif of GMAP-210 (aa 1-38), an artificial coiled-coil and GFP as a fluorescent reporter. The use of the artificial coiled-coil prevents the formation of heterodimers with the GCC construct, hence facilitating the comparison (14, 47). Scale bar: 10 µm.
